# Using single-cell RNA sequencing to generate cell-type-specific split-GAL4 reagents throughout development

**DOI:** 10.1101/2023.02.03.527019

**Authors:** Yu-Chieh David Chen, Yen-Chung Chen, Raghuvanshi Rajesh, Nathalie Shoji, Maisha Jacy, Haluk Lacin, Ted Erclik, Claude Desplan

## Abstract

Cell-type-specific tools facilitate the identification and functional characterization of distinct cell types, which underly the complexity of neuronal circuits. A large collection of existing genetic tools in Drosophila relies on enhancer activity to label different subsets of cells. These enhancer-based GAL4 lines often fail to show a predicable expression pattern to reflect the expression of nearby gene(s), partly due to an incomplete capture of the full gene regulatory elements. While genetic intersectional technique such as the split-GAL4 system further improve cell-type-specificity, it requires significant time and resource to generate and screen through combinations of enhancer expression patterns. In addition, since existing enhancer-based split-GAL4 lines that show cell-type-specific labeling in adult are not necessarily active nor specific in early development, there is a relative lack of tools for the study of neural development. Here, we use an existing single-cell RNA sequencing (scRNAseq) dataset to select gene pairs and provide an efficient pipeline to generate cell-type-specific split-GAL4 lines based on the native genetic regulatory elements. These gene-specific split-GAL4 lines can be generated from a large collection of coding intronic MiMIC/CRIMIC lines either by embryo injection or *in vivo* cassette swapping crosses and/or CRISPR knock-in at the N or C terminal of the gene. We use the developing Drosophila visual system as a model to demonstrate the high prediction power of scRNAseq-guided gene specific split-GAL4 lines in targeting known cell types. The toolkit allows efficient cluster annotation in scRNAseq datasets but also the identification of novel cell types. Lastly, the gene-specific split-GAL4 lines are broadly applicable to Drosophila tissues. Our work opens new avenues for generating cell-type-specific tools for the targeted manipulation of distinct cell types throughout development and represents a valuable resource to the fly research community.

**Significance Statement:** Understanding the functional role of individual cell types in the nervous systems has remained a major challenge for neuroscience researchers, partly due to incomplete identification and characterization of underlying cell types. To study the development of individual cell types and their functional roles in health and disease, experimental access to a specific cell type is often a prerequisite. Here, we establish an experimental pipeline to generate gene-specific split-GAL4 guided by single-cell RNA sequencing datasets. These lines show high accuracy for labeling targeted cell types from early developmental stages to adulthood and can be applied to any tissues in Drosophila. The collection of gene-speicifc-split-GAL4 will provide a valuable resource to the entire fly research community.

---

Proper function of the nervous system relies on the interactions between a large number of different neurons to form circuits. Understanding the tremendous neuronal diversity in the central nervous system and the role of each neuronal type often requires cell-type-specific genetic manipulations. In *Drosophila*, binary expression systems such as GAL4/UAS (1), LexA/LexAop (2), and QF/QUAS (3) systems have been widely used for creating cell-type-specific genetic reagents. For example, a large collection of enhancer-based GAL4 driver lines (e.g., GMR-GAL4 and VT-GAL4) were created, each of which contains 2-3 kb genomic DNA sequences that capture distinct enhancer fragments, allowing genetic access to different subsets of cells (4, 5). Searchable databases for expression patterns in the central brain as well as the ventral nerve cord are available for most of the enhancer-based GAL4 driver lines, facilitating the identification of genetic tools labeling cell types of interest. Although the enhancer-based GAL4 lines labels fewer neurons than enhancer trap GAL4 lines given the use of smaller regulatory genomic sequences, it is often needed to further restrict the expression pattern to achieve desired cell-type specificity by genetic intersectional techniques, such as split-GAL4/LexA systems.

The split-GAL4/LexA system has significantly improved the labeling specificity by splitting GAL4/LexA into GAL4/LexA DNA binding domain (DBD) and transcriptional activation domain (AD), with the expression of each domain under the control of different enhancers (6–9). The genetic intersection is achieved when cells have both enhancers active, thereby reconstituting functional GAL4/LexA expression. While there are thousands of split-GAL4 lines created by using the same enhancer fragments as the original GAL4 lines, presumably having the same expression pattern, these split-GAL4 expression pattern is often inconsistent with the original GAL4 lines (10, 11). This limits the use of the GAL4 expression database for searching specific elements to drive split-GAL4 components. Due to the occasional inconsistency, to generate a highly specific split-GAL4 line for the cell type of interest, one would often need to manually go through the expression image database and look for candidate lines that label the cell type of interest. Subsequently, numerous different combinations of split-GAL4 lines are built and validated experimentally for desired expression pattern. This is often a labor-intensive and time-consuming process. Although the color maximum intensity projection images (color MIPs) algorithm or NeuronBridge software have been recently developed to expedite the search for split-GAL4 combinations (12, 13), it remains challenging to efficiently create cell-type specific split-GAL4 lines with high prediction power.

The time and resource consumption for making cell-type-specific split-GAL4 lines is even more challenging for developmental studies. Many existing cell-type-specific genetic tools are developed for functional studies in the adult central nervous system but often fail to show expression during development. For example, neurotransmitters or their receptors are frequently used as genetic markers for labeling a subset of cell types, but they are not expressed early during development when circuits are forming. The lack of developmental cell-type-specific tools limits the ability to visualize and conduct genetic perturbation for developing neurons. The existing collection of enhancer-based GAL4 or split-GAL4 driver lines often do not reflect the expression of the nearby gene and tend to have random expression patterns, nor do they have systematic documentation of developmental expression patterns, thus hindering the prediction of the expression pattern in a cell-type-specific manner during development.

In this study, we used the developing *Drosophila* visual system as a model to establish a pipeline for generating highly cell-type-specific split-GAL4 lines labeling neurons throughout development. Recent studies have produced single-cell transcriptomic atlases for all neurons at different stages of development of the *Drosophila* visual system and identified around 250 distinct cell types (14, 15). We used these datasets to develop a systematic pipeline for analyzing gene expression in each cell type across various developmental stages. Instead of relying on screening for expression patterns, we identified pairs of genes that are expressed together in only one or very few clusters and generated split-GAL4 drivers that precisely reproduce the expression of the gene at any given developmental stage and together restrict expression to cells where the two genes of the split-GAL4 are co-expressed. Instead of using enhancers that often poorly reflect the expression of the nearby gene, we chose to insert the reporter within the coding sequence using a gene-specific T2A-split-GAL4 approach that much more faithfully reports expression of the gene identified in the developmental scRNAseq. These gene-specific T2A-split-GAL4 lines can be generated through the Recombinase-mediated cassette exchange (RMCE) of T2A-split-GAL4 elements from the large existing collection of MiMIC/CRIMIC lines (16–19) or by N- or C-terminal knock-in of T2A-split-GAL4 elements through CRISPR genome editing. We find high accuracy in labeling predicted cell types using these split-GAL4 lines whose expression was consistent with the scRNAseq expression at the different stages of development. This collection of highly cell-type-specific developmental split-GAL4 lines is a powerful tool to annotate unidentified clusters in scRNAseq datasets and identify novel cell types. Additionally, to expend its applicability, we generated *in vivo* swapping donor flies for creating these gene-specific split-GAL4 lines by genetic crosses, eliminating the need to perform embryo injection. Taken together, the scRNAseq-guided split-GAL4 strategy provides highly cell-type-specific developmental genetic tools to study key genes controlling various neuronal developmental processes. The gene-specific split-GAL4 toolkit developed in our study is adaptable by design to other tissues in *Drosophila* and will be a great resource for the fly community.

## Results

### Pipeline for generating gene-specific T2A-split-GAL4 lines

Our recent scRNAseq dataset of adult Drosophila optic lobe (15) identified about 250 clusters that presumably represent at least 250 different cell types (we will treat these clusters as cell types in the study and assume each cluster is homogenous unless stated otherwise) (Fig. 1A), some of which are categorized into several broad classes such as distal medulla (Dm), lamina (L), lamina wide-field (Lawf), lobula columnar (LC), Lobula Plate intrinsic (LPi), medullar intrinsic (Mi), transmedullary (Tm), and transmedullary Y (TmY) neurons, each of which has distinct morphological features that allow assignment to a specific class (Fig. 1B). We used our scRNAseq data to identify genes that are expressed in a cluster and would thus label only one cell type. However, there is a lack of single and clean marker gene that only labels one cluster in the adult scRNAseq optic lobe dataset, limiting the use of a single GAL4 line reporting the expression of this single gene for cell-type-specific labeling. To broaden the search of cell-type-specific genetic tools for more clusters, we therefore looked for pairs of genes expressed together in one or very few clusters and used the split-GAL4 genetic intersectional strategy where GAL4DBD and GAL4AD (either GAL4AD, p65, or VP16) are under the control of two different sets of genetic regulatory elements (Fig. 1C). We did not use enhancer-based split-GAL4 lines since these enhancer fragments often do not reproduce the expression of the nearby gene and therefore likely do not recapitulate expression of the selected gene. Therefore, we chose the gene-specific-T2A-split-GAL4 approach to better recapitulate the endogenous target gene expression. There are two main ways to create gene-specific-T2A-split-GAL4 lines: 1) T2A-split-GAL4 cassette can be knocked into either the N- or C-terminal of the endogenous gene by CRISPR (Fig. 1D-E); 2) T2A split-GAL4 transgenes can be integrated into coding introns of targeted genes by Recombinase-mediated cassette exchange (RMCE) (Fig. 1F) (16–19). For the latter, a T2A-split-GAL4 donor cassette flanked by attB sequence can be exchanged with a large collection of coding intronic MiMIC/CRIMIC insertions flanked by attP sequence in the presence of C31 integrase. The T2A-split-GAL4 cassette is subsequently spliced into the endogenous RNA transcript via the provided splicing acceptor and donor sequences in the cassette. After translation of the integrated transgenic cassette, the split-GAL4 protein is released from the endogenous truncated protein through a self-cleaving T2A sequence. The expression of these gene-specific-T2A-split-GAL4 should have the same expression pattern as the endogenous target gene since they both use the same native gene regulatory elements.

**Fig. 1.**
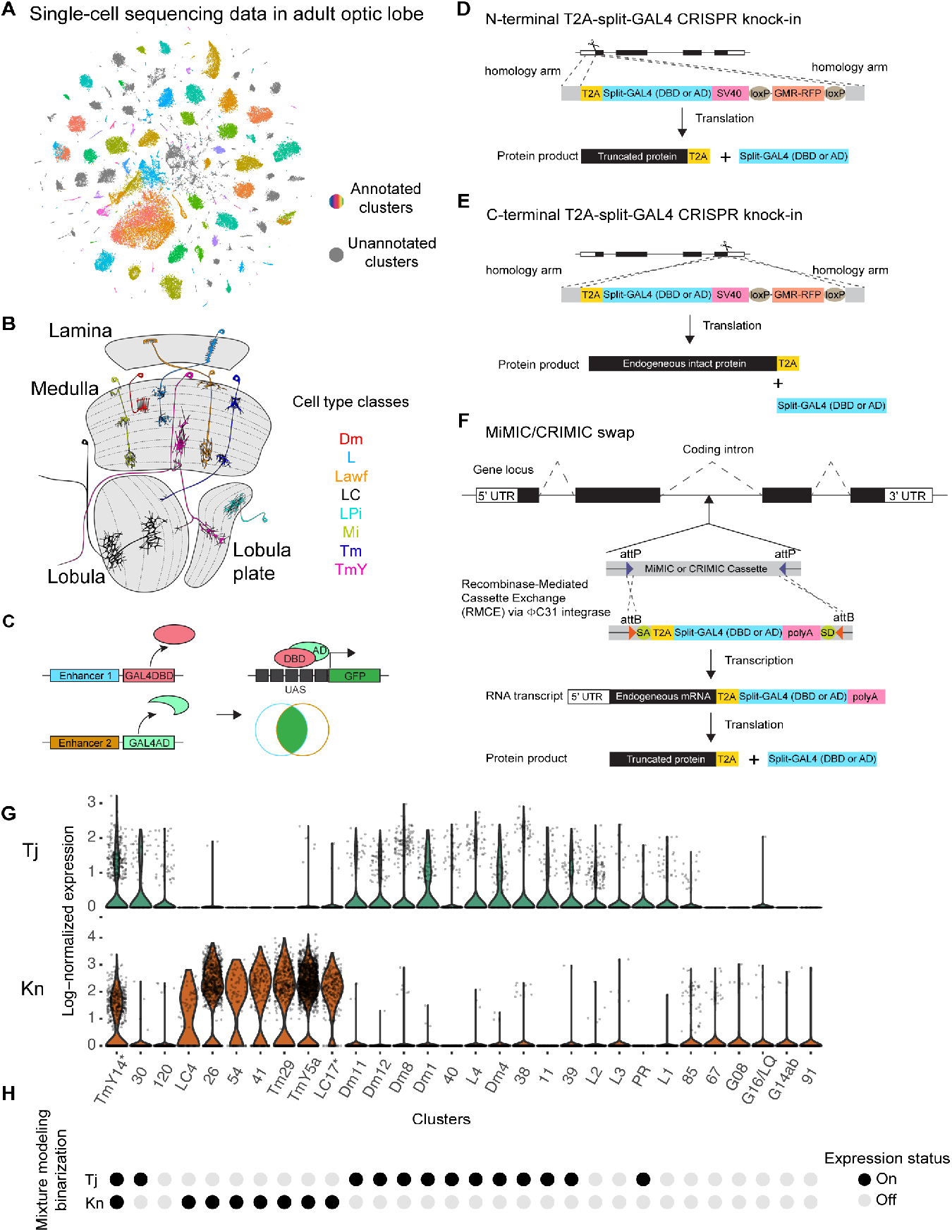
(A) t-SNE plot of the scRNAseq dataset of the adult Drosophila optic lobe. Clusters in gray represent unannotated clusters. Clusters with other colors are annotated in previous study (15). (B) Schematic diagram of the Drosophila visual system with representative cell types in several major classes. There are four main neuropils in the optic lobe: lamina, medulla, lobula, and lobula plate. Distal medulla (Dm), lamina (L), lamina wide-field (Lawf), lobula columnar (LC), Lobula Plate intrinsic (LPi), medullar intrinsic (Mi), transmedullary (Tm), transmedullary Y (TmY). (C) Schematic diagram of the split-GAL4 system. GAL4 can be split into GAL4 DNA-binding domain (GAL4DBD) and activation domain (AD). Labeling specificity is further restricted when GAL4DB and AD are under the control of two different enhancers and only the cells with both enhancers active have a functional reconstituted GAL4 for driving reporter gene expression. (D-F) Schematic diagram of the gene-specific split-GAL4 generation. Gene-specific split-GAL4 lines can be generated either through N- (D) or C-terminal T2A-split-GAL4 knock-in (E), or through Recombinase-mediated cassette exchange (RMCE) of T2A-split-GAL4 elements from the large existing collection of MiMIC/CRIMIC lines (F). The expression of a split-GAL4 reporter depends on the native gene regulatory network and is expected to recapitulate the endogenous transcript expression detected in the scRNAseq dataset. (G) Log-normalized expression of Tj (top) and Kn (bottom) for different clutters. (H) Binarization of gene expression by mixture modeling (15, 20). A probability (ON) score ranging from 0 to 1 is assigned to each cluster: 0 indicates no expression while 1 indicates strong expression. We set the probability score of 0.5 as an expression threshold and classified every gene in every cluster as either ON or OFF. Note that it is practically challenging to define whether a given gene is expressed in each cluster. For example, Tj showed similar expression level between cluster 39 and L2, yet, cluster 39 but not L2 is predicted to express Tj based on mixture modeling binarization.

We next aimed to identify pairs of genes that label each specific cluster of interest. Due to the sparse nature of scRNAseq and the possible presence of ambient RNAs, many droplets would still have zero transcript detected even when a gene is in fact expressed. Ambient RNAs could also introduce low read counts into some droplets even when a gene is not really expressed. It is thus challenging to determine whether a given gene is expressed in a given cluster by a fixed threshold for the expression level at single-cell level (lognorm value) (Fig. 1G). To address this issue, we first binarized the gene expression in the scRNAseq dataset via mixture modeling (15, 20) so that each gene was assigned to an expression probability score [probability (ON)] ranging from 0 to 1 in each cluster, 0 indicating no expression while 1 indicates strong expression. We set the probability score of 0.5 as an expression threshold and classified every gene in every cluster as either ON or OFF (Fig. 1H). For example, after mixture modeling binarization, two transcription factors: Traffic jam (Tj) and Knot (Kn), were predicted to be expressed in 13 and 8 adult clusters, respectively. However, in silico genetic intersection between Tj and Kn predicted to label only one cluster (TmY14) (Fig. 1G-H). The binarization of gene expression in scRNAseq data allows efficient selection of gene pairs for split-GAL4 genetic intersection targeting any given cluster.

### Validation of cell-type specificity for gene-specific T2A-split-GAL4 lines in the adult *Drosophila* optic lobe

We next generated a larger collection of gene-specific T2A-split-GAL4 lines predicted to label various known cell types in the optic lobes. Each cell type tested comprises unique morphological features projecting to specific layer(s) in the neuropil(s). To better ascertain the identity of neurons, we used sparse labeling through the MultiColor FlpOut (MCFO) technique (21). We validated our in silico prediction by crossing these split-GAL4 lines with a UAS-GFP reporter for full expression or to MCFO lines for sparse labeling. Five split-GAL4 combinations were predicted to be only present together in one cluster and thus should specifically label only one cell type: TmY14 was labeled by Tj-DBD + Kn-p65 (Fig. 2A), Dm1 was labeled by CG5160-DBD + DIP-GAL4AD (Fig. 2B), Dm12 was labeled by Tj-DBD + Dop1R2-VP16 (Fig. 2C), Mi15 was labeled by CngA-DBD + Ple-p65 (Fig. 2D), and LPLC2 was labeled by Acj6-DBD + DIP -VP16 (Fig. 2E). There are more than one split-GAL4 combinations labeling the same cluster. For further validation of this approach, we asked whether we could observe the targeted cluster by using different combinations of genes. We found that Dm1 could also be labeled by CG5160-DBD + Tj-VP16 (in addition to CG5160-DBD + DIP-GAL4AD used above). We tested the expression of this split-GAL4 line and found that it indeed label Dm1, although sparse labeling confirms the presence of additional neurons, such as Dm11 and L2 (SI Appendix, Fig. S1A).

**Fig. 2.**
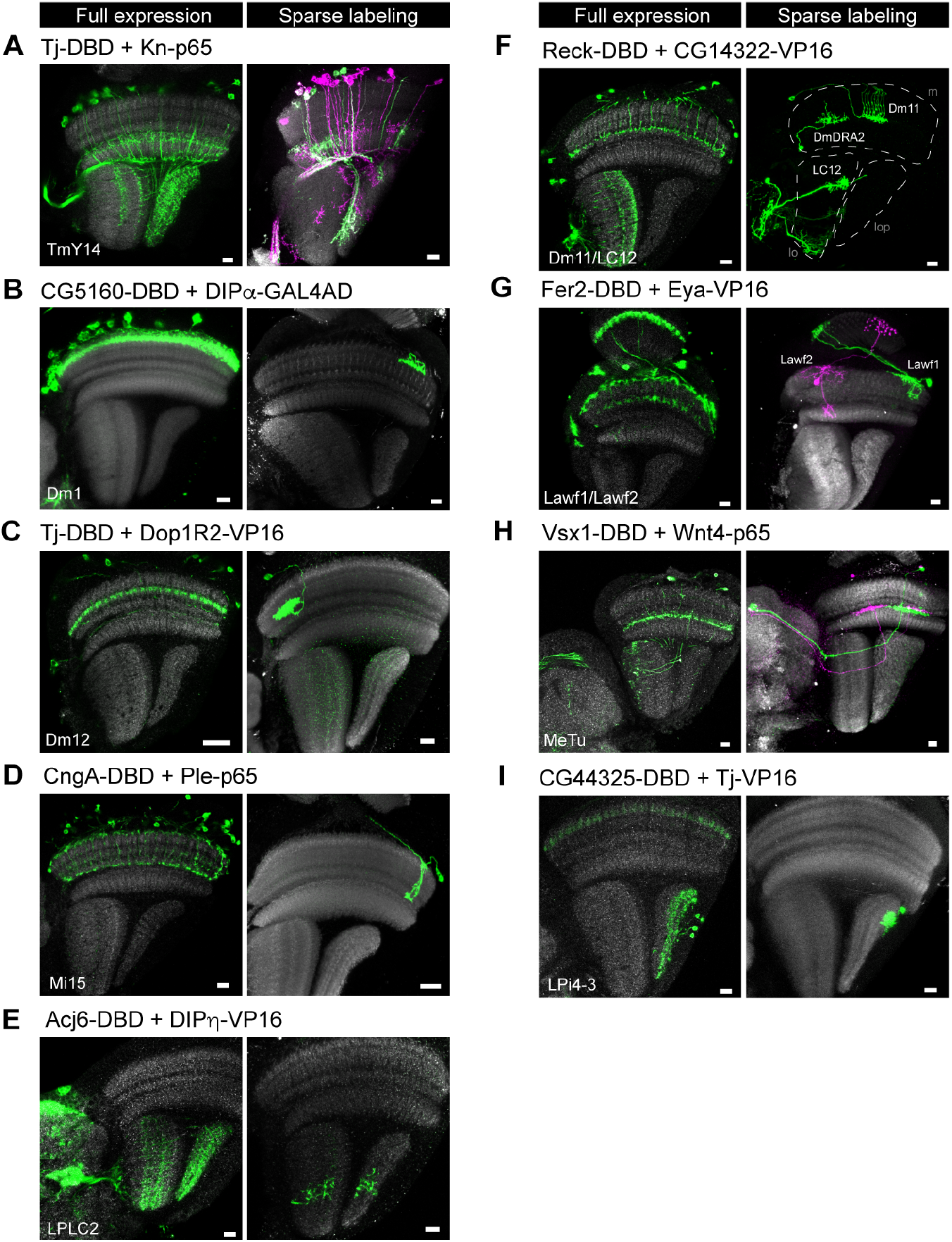
Characterization of selected gene-specific split-GAL4 lines targeting different cell types/clusters. Targeted cell types predicted by scRNAseq expression are shown in the lower left corner for each split-GAL4 line. The expression pattern of each split-GAL4 line is shown either with UAS-myr-GFP reporter for full expression (right) or with UAS-MCFO lines for sparse labeling (right). (A-E) Examples of split-GAL4 lines targeting single cell types. (F-G) Example of split-GAL4 lines targeting two cell types. (H-I) Examples of split-GAL4 lines targeting unannotated cell types in the scRNAseq dataset. Anti-NCad staining (gray) is used for visualizing neuropils. Images are substack projections from full expression labeling or segmented single cells from sparse labeling to show distinct morphological features of distinct cell types. Scale bar: 10 μm.

We next used two lines that were predicted to each label two known clusters: Dm11 and LC12 were both labeled by Reck-DBD + CG14322-VP16 (Fig. 2F) while Lawf1 and Lawf2 were both labeled by Fer2-DBD + Eya-VP16 (Fig. 2G). Another distinct combination of genes was predicted to label Dm11 and LC12: Indeed CG9896-GAL4DBD + CG14322-VP16 labeled these two cell types (SI Appendix, Fig. S1B). Additional cell types were also labeled by this combination, such as the neurons from cluster 30 (named MeSps neurons, see later description) and another unknown neuron in the lobula (SI Appendix, Fig. S1B-C). Our results showed that all target cell types were labeled by the predicted split-GAL4 combinations, although additional cell types might be observed in some cases.

### Gene-specific T2A-split-GAL4 line uncovers a novel cell type

The high accuracy in predicted clusters using the gene-specific T2A-split-GAL4 lines facilitates not only assigning known cell types to unannotated clusters in our scRNAseq data (Fig. 2H-I), but also identifying novel cell types that have not been described. In our scRNAseq dataset, we found that TkR86C and CG14322 is expressed in 12 and 8 clusters, respectively (Fig. 3A). However, cluster 30, an unannotated cluster, is the only cluster after in silico genetic intersection of these two genes (Fig. 3A). We generated a gene-specific split-GAL4 line with (TkR86C-DBD + CG14322-VP16) and examined its expression in the adult optic lobe. We found TkR86C-DBD + CG14322-VP16 split-GAL4 line labeled a novel type of medulla projection neuron whose cell body is located in the medulla cortex and its neurites innervate medulla layer M7 (the serpentine layer) (Fig. 3B-C). Sparse labeling revealed that cluster 30 neurons bifurcate at M7 layer with one neurite branch innervating multiple visual columns (Fig. 3D-E) and another branch running along the border of the medulla and lobula and projecting to the superior posterior slope (Sps) in the central brain (Fig. 3F-G). The bifurcation of neurites in the medulla can be bidirectional (Fig. 3E) and all medulla columns are innervated by a total of 40-50 cells (Fig. 3B-C). We named this novel medulla projection neuron Medulla-Superior posterior slope, MeSps.

**Fig. 3.**
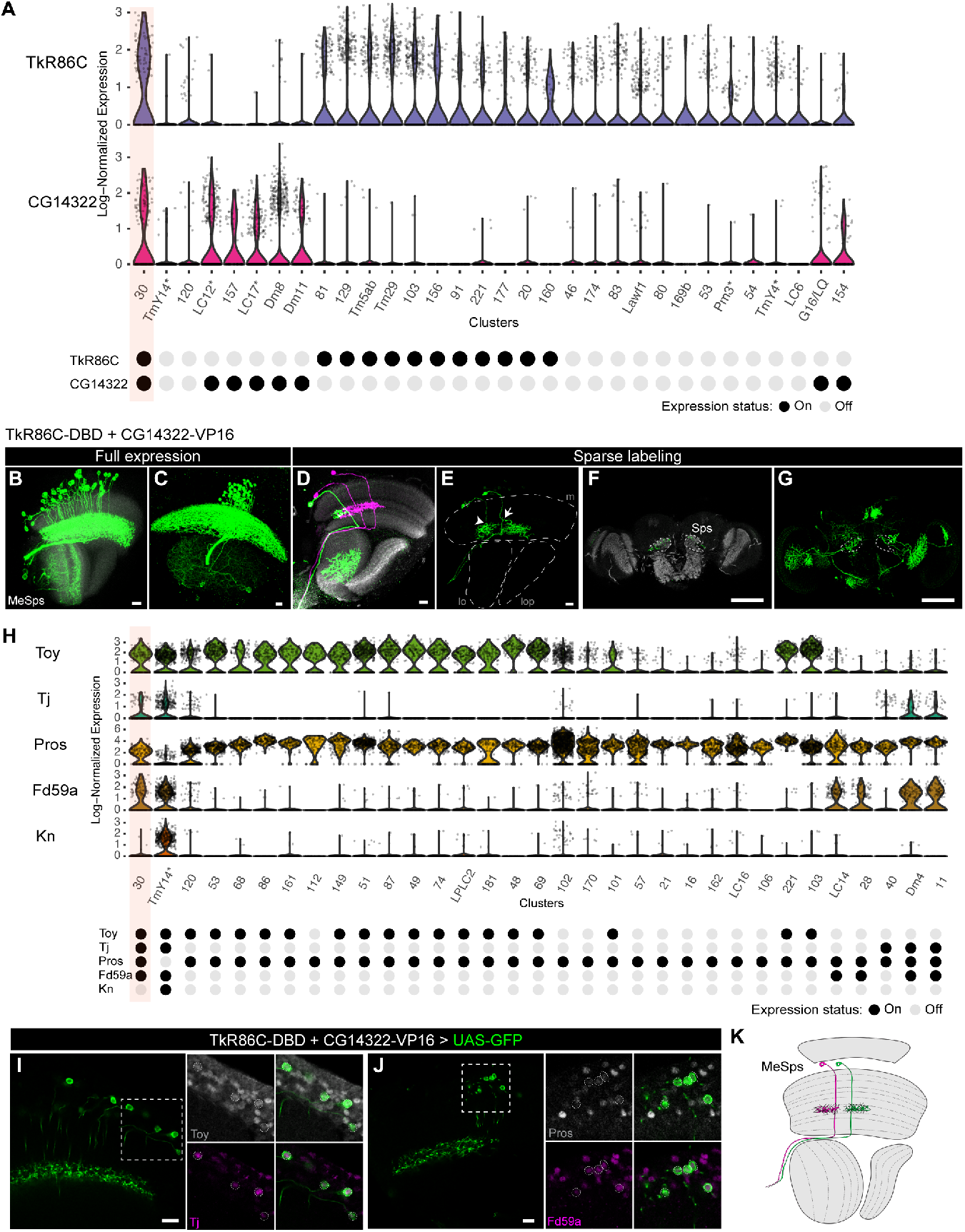
(A) Log-normalized expression of TkR86C and CG14322 for different clutters (Top). Mixture modeling binarization of expression status for both genes is shown at the bottom. Note that cluster 30 is predicted to be the only cluster intersected by TkR86C and CG14322. (B-C) The full expression pattern of TkR86C-DBD + CG14322-VP16 line is shown with UAS-myr-GFP reporter. Note that there is an additional cell type in the lobula with only 2-3 cells labeled. The cell bodies of MeSps neurons are restricted to the medulla cortex. (D-E) Sparse labeling of MeSps neurons using MCFO. The bifurcation of neurites at M7 layer can be bidirectional (arrowhead to the anterior projection; arrow to the posterior projection). m: medulla; lo: lobula; lop: lobula plate. Scale bar: 10 m. (F-G) Sparse labeling of MeSps neurons using MCFO show their projection to the superior posterior slope (Sps). Single optical section (F) or substack maximum projections (G) are shown. Scale bar: 100 m. Anti-NCad staining (gray) is used for visualizing neuropils. (H) Log-normalized expression of selected transcription factors (Toy, Tj, Pros, Fd59a, and Kn) (Top). Mixture modeling binarization of expression status for selected genes are shown at the bottom. Note that cluster 30 is predicted to be the only cluster positive for Toy, Tj, Pros, and Fd59a. Kn is used to serve as a negative marker for cluster 30. (I-J) Co-staining of MeSps neurons labeled by TkR86C-DBD and CG14322-VP16 with anti-Toy (Gray) and anti-Tj (Magenta) in I and anti-Pros (Gray) and anti-Fd59a (Magenta) in J. MeSps neurons expressing GFP reporter are outlined in dotted circles. Scale bar: 10 μm. (K) Schematic diagram of MeSps neurons, a novel type of medulla projection neurons.

Since MeSps neurons have never been described before, we performed immunofluorescence staining for additional transcription factor markers that collectively define cluster 30 in the scRNAseq data (Fig. 3H). This confirmed that MeSps neurons are positive for Toy, Tj, Pros, and Fd59a (Fig. 3I-J), while negative for Kn (SI Appendix, Fig. S2). In conclusion, we identified a novel medulla projection neuron by using the gene-specific split-GAL4 approach (Fig. 3K), and we have high confidence in assigning cluster 30 to the MeSps neurons labeled by this split-GAL4 line due to the additional staining results of four positive transcription factor markers.

### Gene-specific T2A-split-GAL4 lines label neurons from early developmental stages to adulthood

There are many existing split-GAL4 lines generated by the enhancer-based approach that specifically label various cell types in the Drosophila optic lobe (20, 22, 23). However, most of the lines do not label the same cell types during pupal development, and some of the drivers are expressed in different neuronal types during development. Taking advantage of the high prediction accuracy of our scRNAseq-guided gene-specific split-GAL4 approach and the availability of single-cell transcriptomes of all optic lobe neurons in the optic lobe at five pupal stages (P15, P30, P40, P50, and P70), we selected split-GAL4 lines where both genes were co-expressed at most if not all pupal stages. We tested four different split-GAL4 combinations targeting the TmY14 (Fig. 4A), MeSps (Fig. 4B), Dm11/LC12 (Fig. 4C), and LPi4-3 (Fig. 4D) and examined the expression at two earlier developmental stages (P15 and P50). All four split-GAL4 drivers consistently labeled the predicted cell types at these time points, albeit additional cell types were observed (Fig. 4A-D). For example, a lobula columnar neuron was labeled by the MeSps split-GAL4 driver (Fig. 4B); lamina neurons were labeled in LPi4-3 split-GAL4 driver (Fig. 4D). Another split-GAL4 combination (Beat-IIIc-GAL4DBD + DIP-GAL4AD) predicted to label LPi4-3 from P15 to P70 did show LPi4-3 labeling exclusively at P15 but showed at least two additional cell types in adults (one in lobula and another in medulla) (SI Appendix, Fig. S3). These results support the consistency of predicted cell types labeled across developmental stages by the scRNAseq-guided gene-specific split-GAL4 approach.

**Fig. 4.**
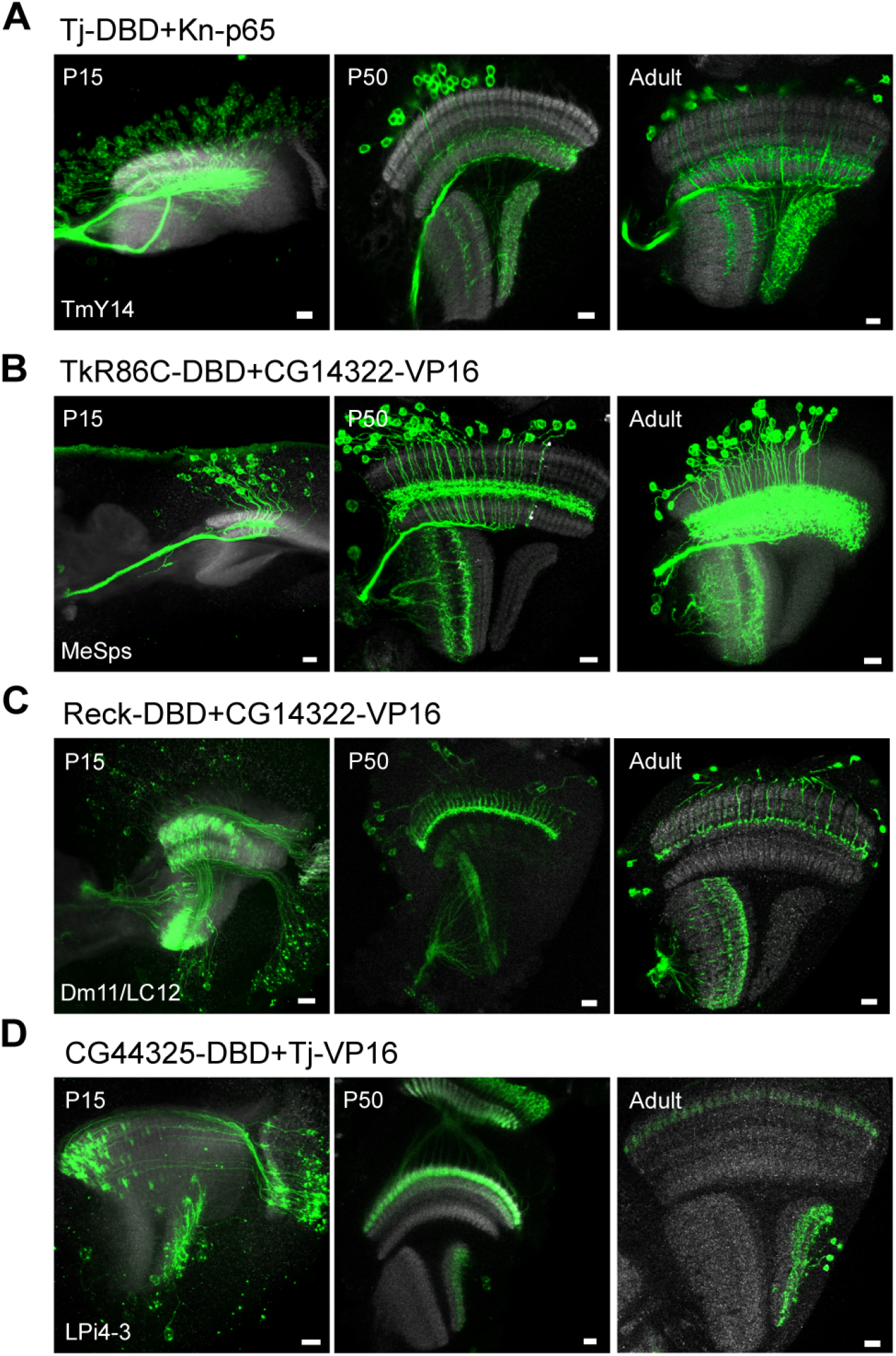
Developmental characterization of selected gene-specific split-GAL4 lines targeting different cell types/clusters. Targeted cell types predicted by scRNAseq expression are shown in the lower left corner for each split-GAL4 line. The full expression pattern of each split-GAL4 line is shown with UAS-myr-GFP reporter at P15, P50, and adult stages. Note that the targeted cell types are always observed at multiple developmental stages, although other cell types might be observed. Anti-NCad staining (gray) is used for visualizing neuropils. Images are substack projections from full expression labeling to show distinct morphological features of distinct cell types. Scale bar: 10 *μ*m.

### Extensive predicted optic lobe cluster coverage by marker gene pairs

There are around 250 clusters in our recent single-cell developmental transcriptomic atlas (15). After merging developmental subclusters and excluding immature clusters, ganglion mother cells (GMCs), or transient extrinsic (TE) neurons that are only present during development and not in the adult, there are 198 clusters having a corresponding cluster in the adult. We aimed to determine how many clusters could be labeled by combinations of gene pairs throughout development using the scRNAseq-guided gene-specific split-GAL4 approach (Fig. 5). We first defined an ideal gene pair as the one that labels only a single cluster for all six developmental stages while none of the other clusters showed any expression by this gene pair at any developmental stages (Fig. 5A). We found 91 clusters that can be labeled by ideal gene pairs and cell-type-specific split-GAL4 driver lines targeting these clusters could theoretically be generated. When allowing expression in 1 or 2 additional non-ideal clusters (defined as the clusters that show expression at most 2 developmental stages, either the same non-ideal cluster expressing the gene pair at 2 developmental stages or two non-ideal clusters expressing the gene pair at 1 developmental stage each), 33 additional clusters could be labeled (Fig. 5D). These data suggest that 62.6% of the clusters (124/198) can be labeled by split-GAL4 gene pairs that can be used as developmental drivers since both genes are predicted to be expressed from P15 to adult.

**Fig. 5.**
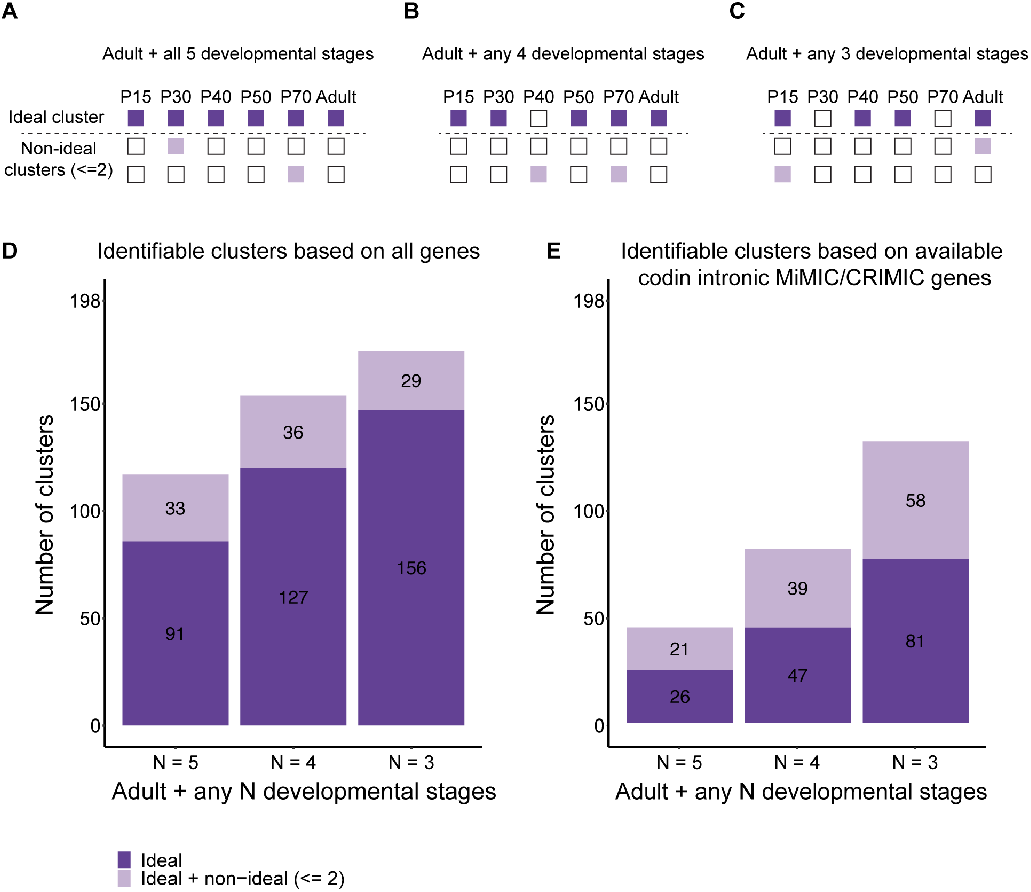
(A-C) Illustration of strategies to identify gene pairs that mark a cluster of interest throughout development with different stringency. Dark violet boxes indicate the stages when a gene pair is predicted to be on in the cluster of interest while light violet boxes indicate when the gene pair is predicted to be active in other clusters. (D) Number of clusters that are predicted to be identified with gene pairs suggested by each strategy when all genes detected in the atlas are considered. (E) Number of clusters that are predicted to be identified with gene pairs suggested by each strategy when only considering genes with coding intronic MiMIC or CRIMIC lines available for recombinase-mediated cassette exchange (3637 genes).

A broader search of potential gene pairs can be done by relaxing the definition of the ideal clusters as a cluster expressing a given gene pair at adult plus several developmental stages. The more relaxed search strategy addresses the scenario when a gene combination is only predicted to be not active in a minority of developmental stages. While this could well reflect biological variations, considering the sensitive nature of the binary expression system, it is likely that split-GAL4 perdures during these stages, and the driver line would still behave as active throughout all the stages despite transcriptional fluctuation. We always require the ideal cluster to have the adult stage labeled to ensure unambiguous identification of neuronal morphology. When we only look for expression at adult plus any four out of five (Fig. 5B) or any three out of five developmental stages (Fig. 5C) for the ideal cluster, then most clusters are covered by a pair of genes (127/198, 64% for ideal gene pairs, 163/198, 82.3% for including ideal and non-ideal gene pairs when selecting adult plus any four out of five developmental stages; 156/198, 78.8% for ideal gene pairs, 185/198, 93.4% for including ideal and non-ideal gene pairs when selecting adult plus any three out of five developmental stages). Each ideal cluster can often be labeled by more than one combination of gene pairs, and we listed all possible combinations of gene pairs that are expressed at all six developmental stages in SI Appendix, Table. S3.

Given that there are many genes with readily split-GAL4 convertible coding intronic MiMIC or CRIMIC lines, we also examined the number of clusters with ideal gene pairs currently available for cost-efficient split-GAL4 conversion (Fig. 5E). We found 47, 86, 149 out of 198 clusters that have both ideal and non-ideal gene pairs with available coding intronic MiMIC or CRIMIC for adult plus five, four, and three developmental stages, respectively (Fig. 5E).

Annotation of scRNAseq clusters is challenging and there are still 112 out of 198 clusters in the optic lobe that remain unannotated in the optic lobe developmental atlases. To examine the number of unannotated clusters that can be covered using our approach, we performed similar analyses with the 112 unannotated clusters. We found 95.6% cluster coverage (107/112) when using gene pairs targeting both ideal and non-ideal clusters in adult plus three other developmental stages (SI Appendix, Fig. S4). These results suggest that combinations of genes can be found to generate split-GAL4 lines for most if not all the clusters in the optic lobe and more than half of the clusters have readily convertible split-GAL4 reagents.

### *in vivo* swapping for generating gene-specific T2A-split-GAL4 lines from the collection of coding intronic MiMIC/CRIMIC

One of the features of gene-specific T2A-split-GAL4 lines is the adaptability for other tissues in Drosophila as long as the target gene is expressed in the tissue of interest. There is a large collection of coding intronic MiMIC/CRIMIC lines deposited in the Bloomington Drosophila Stock Center that can be readily swapped into either GAL4DBD or AD split-GAL4 drivers via embryo injection of an appropriate donor DNA template (17). We adapted the *in vivo* swapping method for making T2A-GAL4 lines (17) and modified it for making T2A-split-GAL4 lines (Fig. 6A). Briefly, the T2A-split-GAL4 cassettes with reading frames 0, 1, and 2 were flanked by three lox sequence variants where a circular donor can be excised in the presence of Cre recombinase. By providing both Cre recombinase and C31 integrase, the T2A-split-GAL4 cassette with the right orientation (forward or reverse) and correct splicing phase (phase 0, 1, or 2) can be selected by genotyping PCR. We generated triple donor flies that carry donor cassettes for all three splicing phases and tested the *in vivo* swapping by using the bru1[MI00135] coding intronic MiMIC line (SI Appendix, Fig. S5). bru1 is encoded in the + strand and the bru1[MI00135] is in phase 1 (Fig. 6B); therefore, the successful swap that generates bru1-split-GAL4 should have a forward integration with phase 1 donor.

**Fig. 6.**
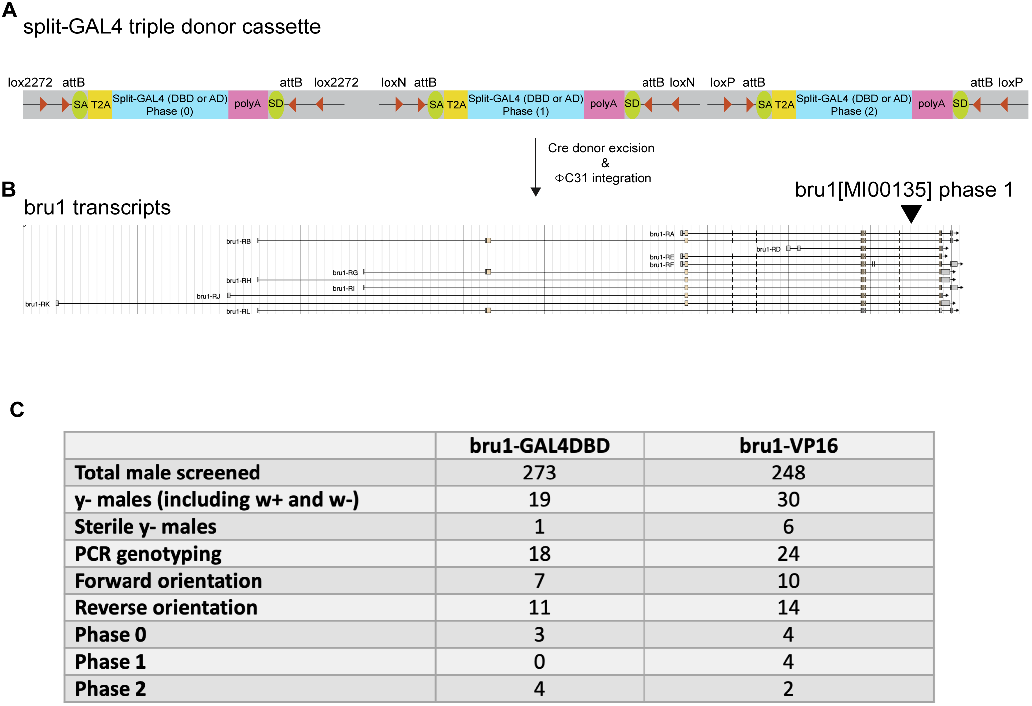
(A) Schematic diagram showing T2A-split-GAL4 triple donor cassette. The cassette is modified from T2A-GAL4 triple donor (Diao et al., 2015) by replacing T2A-GAL4 with T2A-GAL4DBD or T2A-VP16AD. Split-GAL4 donors with three different splicing phases (phase 0, 1, and 2) are flanked by attB sequences and lox sequences variants. Supplement of Cre recombinase and C31 integrase will allow integration of T2A-split-GAL4 into targeted MiMIC insertion. (B) Gene structure annotation of bru1 (encoded in a + orientation) from JBrowse using D. melanogaster (r6.49) reference (24). Phase 1 coding intronic MiMIC insertion (MI00135) is highlighted by an arrowhead. (C) Summary table of *in vivo* swapping by triple T2A-split-GAL4 donor. The genetic crossing scheme is shown in SI Appendix, Fig. S4.

We performed GAL4DBD, and VP16 triple donor crosses with bru1[MI00135] coding intronic MiMIC line and selected 273 and 248 y- males for GAL4DBD and VP16 lines for further molecular mapping, respectively. We expected to have 50/50 of the forward and reverse cassette integration events when selecting for y- (MiMIC) flies. For each orientation, there should be one-third of the flies for each phase. We observed 38.8% (forward orientation) and 61.2% (reverse orientation) for GAL4DBD swapping, 41.6% (forward orientation), and 58.4% (reverse orientation) for VP16 swapping (Fig. 6C). We performed PCR genotyping for the individuals with forward integration and identified no individuals with phase 1 for GAL4DBD donors and 4 individuals with phase 1for VP16 donors (Fig. 6C). Although we did not obtain bru1-GAL4DBD with the correct phase, we successfully obtained bru1-VP16 lines with the correct integration orientation and splicing phase. While that the triple donor simplifies the crossing scheme when performing a large-scale *in vivo* swapping, the simplicity came at a cost of reducing the success split-GAL4 line to one-third. To provide a versatile platform for *in vivo* swapping, we also generated single donor split-GAL4 lines to increase the success rate of correct cassette integration to 50%. In sum, we generated both triple donors and single donor T2A-split-GAL4 flies for *in vivo* swapping and provided a proof-of-principle example with a successful swapping of bru1-VP16. These tools will be valuable for the fly community to generate gene-specific T2A-split-GAL4 lines at a low cost.

## Discussion

Having a cell-type-specific genetic tool is often the prerequisite for many genetic manipulations used to study development and function of neurons. Using the Drosophila visual system as a model for which a developmental transcriptomic atlases exists (14, 15), we showed that scRNAseq-assisted gene-specific split-GAL4 lines provide highly predictable and specific labeling for most cell types of interest. As compared to the conventional screening of enhancer-based split-GAL4, this gene-specific split-GAL4 offers the following advantages:

1. Gene-specific split-GAL4 lines have high predictive accuracy. For all 14 gene-specific split-GAL4 lines tested in this study, we always observed that the expression of these split-GAL4 lines was the predicted cell types. This facilitates the annotation of unknown clusters in scRNAseq datasets. We also took advantage of the high prediction accuracy to select gene pairs that are also expressed throughout most, if not all, developmental stages in the developing optic lobe. Our expression analyses showed that these split-GAL4 drivers indeed label the targeted cell types during development and are therefore very useful developmental drivers. Although additional cell types are sometimes observed. these cell types might represent rare cell types that do not have a cluster in our scRNAseq clustering. Alternatively, there might be a false negative in our mixture modeling. Having additional cell types in the split-GAL4 lines might provide genetic access to these cell types that might be of interest.
2. The prediction of the recombination of two split-GAL4 hemi drivers is made possible since location of each gene is known. The large collection of enhancer-based split-GAL4 lines exists either inserted at the attP2 and/or attP40 sites, thereby limiting the possible recombination when both GAL4DBD and AD are inserted at the same locations. For example, all the VT-GAL4DBD lines were inserted at attP2 while one-third of VT-AD lines were also inserted at attP40 (11). There are also concerns about the potential side effects of these attP sites (25–27), creating further complications when using these enhancer-based split-GAL4 lines. In contrast, two gene-specific split-GAL4 could be recombined when present on the same chromosome and their locations could predict the recombination frequency (e.g., TkR86C-DBD + CG14322-VP16 for MeSps and Reck-DBD + CG14322-VP16 for Dm11/LC12 in this study).
3. There are multiple split-GAL4 combinations that are predicted to target the same cluster, providing independent drivers for genetic manipulations of the same cell type. For example, both Reck-DBD + CG14322-VP16 and CG9896-DBD + CG14322-VP16 can be used to manipulate Dm11 and LC12 neurons. The former can be recombined on a single chromosome, providing further flexibility for researchers to choose the desired combinations of split-GAL4. We noted that coding intronic MiMIC/CRIMIC-derived split-GAL4 create mutant alleles for the targeted gene. Performing experiments in a heterozygous background should alleviate this issue and MiMIC/CRIMIC-T2A-GAL4 lines are indeed widely used. Nevertheless, proper controls are necessary, including performing experiments with another combination of split-GAL4 targeting the same cell type.
4. The gene-specific split-GAL4 lines are adaptable to other tissues in Drosophila. With the development of scRNAseq technology, single-cell transcriptomes for various tissues are accumulating rapidly and expanding the list of tissues that are applicable to generate split-GAL4 lines for genetic manipulation. Our pipeline in binarizing scRNAseq expression matrix using mixture modeling can be readily applied to other systems with existing scRNAseq data. These tools would further simplify the search for desired split-GAL4 combination.
5. Thousands of gene-specific split-GAL4 combinations can be readily generated. There are currently available coding intronic MiMIC/CRIMIC lines corresponding to 3637 genes, and additional new CRIMIC lines are still being generated. The gene-specific split-GAL4 lines will become a powerful resource once all these lines are converted to split-GAL4 lines. The triple and single donor split-GAL4 lines generated in this study would be perfectly suited for a large-scale in vivo swapping of these coding intronic MiMIC/CRIMIC lines into split-GAL4 lines simply by crosses. For the genes without coding intronic MiMIC/CRIMIC lines, the advanced CRISPR genome editing methods can perform T2A-split-GAL4 knock into essentially any target.

In sum, we developed an scRNAseq-guided approach for generating highly predictable cell-type-specific T2A-split-GAL4 lines that can be adaptable to any tissue in Drosophila. These genetic reagents enable genetic access for labeling specific neuronal populations in the fly throughout development and will be a valuable resource for understanding the regulation of gene networks that confer cellular morphology, physiology, function, and identity.

## Materials and Methods

### Fly strains

Flies were reared on molasses-cornmeal-agar food at 25°C. The fly lines used in the paper are described in SI Appendix, Table. S1. CRISPR-mediated T2A-split-GAL4 knock-in was performed by WellGenetics Inc. (Taipei, Taiwan). In brief, the gRNA sequences (specified in SI Appendix, Table. S2) were cloned into U6 promoter plasmid(s). Cassette T2A-GAL4DBD (or VP16AD)-GMR-RFP, which contains T2A, Zip-GAL4DBD (or VP16AD), SV40 3UTR, a floxed GMR-RFP, and two homology arms were cloned into pUC57-Kan as donor template for repair. Targeting gRNAs and hs-Cas9 were supplied in DNA plasmids, together with donor plasmid for microinjection into embryos of control strain w1118. F1 flies carrying selection marker of GMR-RFP were further validated by genomic PCR and sequencing.

Split-GAL4 lines generated by injecting donor DNA plasmids into F1 embryos from crosses of coding intronic MiMIC and CRIMIC lines with C31 integrase source. Embryo injections for transgenesis and transformant recovery was completed by BestGene (Chino Hills, CA). The split-GAL4 donor plasmids with appropriate splicing phases were obtained from Addgene, including pBS-KS-attB2-SA(0)-T2A-GAL4DBD-Hsp70 (Addgene #62902), pBS-KS-attB2-SA(1)-T2A-GAL4DBD-Hsp70 (Addgene #62903), pBS-KS-attB2-SA(2)-T2A-GAL4DBD-Hsp70 (Addgene #62904), pBS-KS-attB2-SA(0)-T2A-VP16AD-Hsp70 (Addgene #62905), pBS-KS-attB2-SA(1)-T2A-VP16AD-Hsp70 (Addgene #62908), and pBS-KS-attB2-SA(2)-T2A-p65AD-Hsp70 (Addgene #62915). The integration orientation of yellow progenies from MiMIC injection and 3xP3-GFP-progenies from CRIMIC injection were confirmed by PCR genotyping. See also SI Appendix, Table. S2 for further information.

### Triple and single split-GAL4 donor transgenic lines

For generating triple donor GAL4DBD plasmid pC(lox2-attB2-SA-T2A-GAL4DBD-Hsp70)3, we synthesized SphI_T2A-GAL4DBD_NotI, MluI-T2A-GAL4DBD_FesI, and BsiWI-T2A-GAL4DBD_AscI by GenScript (Piscataway, USA). The three fragments were subcloned into pC-(lox2-attB2-SA-T2A-Gal4-Hsp70)3 (Addgene #62957) by replacing T2A-Gal4 with T2A-GAL4DBD. For generating triple donor VP16 plasmid pC(lox2-attB2-SA-T2A-VP16-Hsp70)3, we synthesized SphI_T2A-VP16_NotI, MluI-T2A-VP16_FesI, and BsiWI-T2A-VP16_AscI by GenScript (Piscataway, USA). The three fragments were subcloned into pC-(lox2-attB2-SA-T2A-Gal4-Hsp70)3 (Addgene #62957) by replacing T2A-Gal4 with T2A-VP16. Both triple donor plasmids were sent for standard P-element-mediated transformation performed by WellGenetics Inc. (Taipei, Taiwan).

For single donor split-GAL4 plasmids, we synthesized BamHI_T2A-GAL4DBD_BamHI and BamHI_T2A-VP16_BamHI by GenScript (Piscataway, USA). The fragments were subcloned into either pC-(loxP2-attB2-SA(0)-T2A-Gal4-Hsp70), Addgene #62954, pC-(loxP2-attB2-SA(1)-T2A-Gal4-Hsp70), Addgene #62955, or pC-(loxP2-attB2-SA(2)-T2A-Gal4-Hsp70); Addgene #62956 by replacing T2A-Gal4 with T2A-GAL4DBD or T2A-VP16. The single donor plasmids were sent for standard P-element-mediated transformation performed by BestGene (Chino Hills, CA).

### Immunohistochemistry

Flies were anesthetized on ice, and the optic lobes were dissected in ice-cold Schneider’s Drosophila Medium (Thermo Fisher, #21720024) for less than 30 min. Tissues were fixed for 20 min with 4% paraformaldehyde in 1X Dulbecco’s phosphate-buffered saline (DPBS, Corning, #21031CV) at room temperature. After three washes with 1X PBST (DPBS with 0.3% Triton X-100), tissues were incubated in primary antibody solutions for 2 days at 4°C. Samples were washed with 1X PBST for at least 6 times (15 min per wash) followed by incubating in secondary antibodies 1-2 days at 4oC. Samples were washed again with 1X PBST for at least 6 times (15 min per wash) followed by the last wash in 1X DPBS. Both primary and secondary antibodies were prepared in 1X PBST with 5% goat serum (Thermo Fisher, #16210064). The antibodies used in the paper are described in SI Appendix, Table. S1. Samples were mounted in VECTASHIELD antifade mounting medium (Vector Laboratories, #H-1000) and stored at 4°C. Fluorescent images were acquired using a Leica SP8 confocal microscope with 400 Hz scan speed in 1024×1024 pixel formats. Image stacks were acquired at 0.5-1 m optical sections. Unless otherwise noted, all images were presented as maximum projections of the z stack generated using Leica LAS AF software.

### Identification of marker gene pairs with mixture modeling-inferred binarized expression

To identify marker gene pairs (two genes) that are specific to a cluster, we implemented a greedy search algorithm to minimize the number of clusters that express a given gene pair. Briefly, for each cluster, we begin with a gene that (1) is expressed in the cluster of interest (target cluster) and (2) is expressed in fewest other clusters (off-target cluster). The same steps are repeated once only among the clusters positive for the first gene selected to identify split-GAL4 candidates but can potentially be extended to identify combinations consisting of more than two genes.

To determine whether a gene is expressed in a cluster, we assign probability of whether a gene is expressed (*P* (*ON*)) to each cluster at each stage as previously described (15, 20). Briefly, we define (1) a baseline unimodal model that cluster average expression of a given gene follows a Gaussian distribution and (2) an alternative bimodal model that cluster average expression follows a mixture of two Gaussian distributions, representing ON and OFF respectively. Parameters of the two models were estimated, and the wellness of fit were compared with expected log predictive density (elpd, a measure of how well a model explains observed data) in 8-fold cross-validation in Stan (28), a software package that implements Bayesian probabilistic model fitting with Markov chain Monte Carlo algorithm. Genes that fit better with the bimodal model 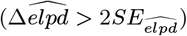 were considered as bimodally distributed while other genes were considered as unimodal. Probability of whether a gene is expressed [*P* (*ON*)] in a cluster were then estimated with the models fitted above with R 4.0.4 (29) and mclust (30).

Gene x cluster probability matrices were generated for datasets corresponding to each stage, and we consider a gene with *P* (*ON*) > 0.5 as expressed. This process is equivalent to adaptively determining gene-specific binarization thresholds with Gaussian mixture models. Binary expression states not only allow efficient greedy combination search described above, but also enabled versatile integration across time series via Boolean algebra. Reduced computation resource requirement with binary data also allowed comprehensive marker combination search.

## Supporting information

Link to the Materials and Methods

Link to the Materials and Methods

Link to the Result

## ACKNOWLEDGMENTS

We thank Fengqiu Diao and Ben White for providing technical assistance on designing split-GAL4 triple donor constructs. We thank all the Desplan lab member for critical feedback on this work, especially Isabel Holguera, Bogdan Sieriebriennikov, Ryan Loker and Sromana Mukherjee. Stocks were also obtained from the Bloomington Drosophila Stock Center (NIH P40OD018537). This work was funded by grants from the NIH (5R01EY013010 to C.D.). Y.-C.D.C. is supported by NIH NRSA (5F32EY032750). Y.C.C. was supported by the MacCracken Program at New York University, by a NYSTEM institutional training grant (Contract #C322560GG), and by Scholarship to Study Abroad from the Ministry of Education, Taiwan.

---

**Fig. S1.**
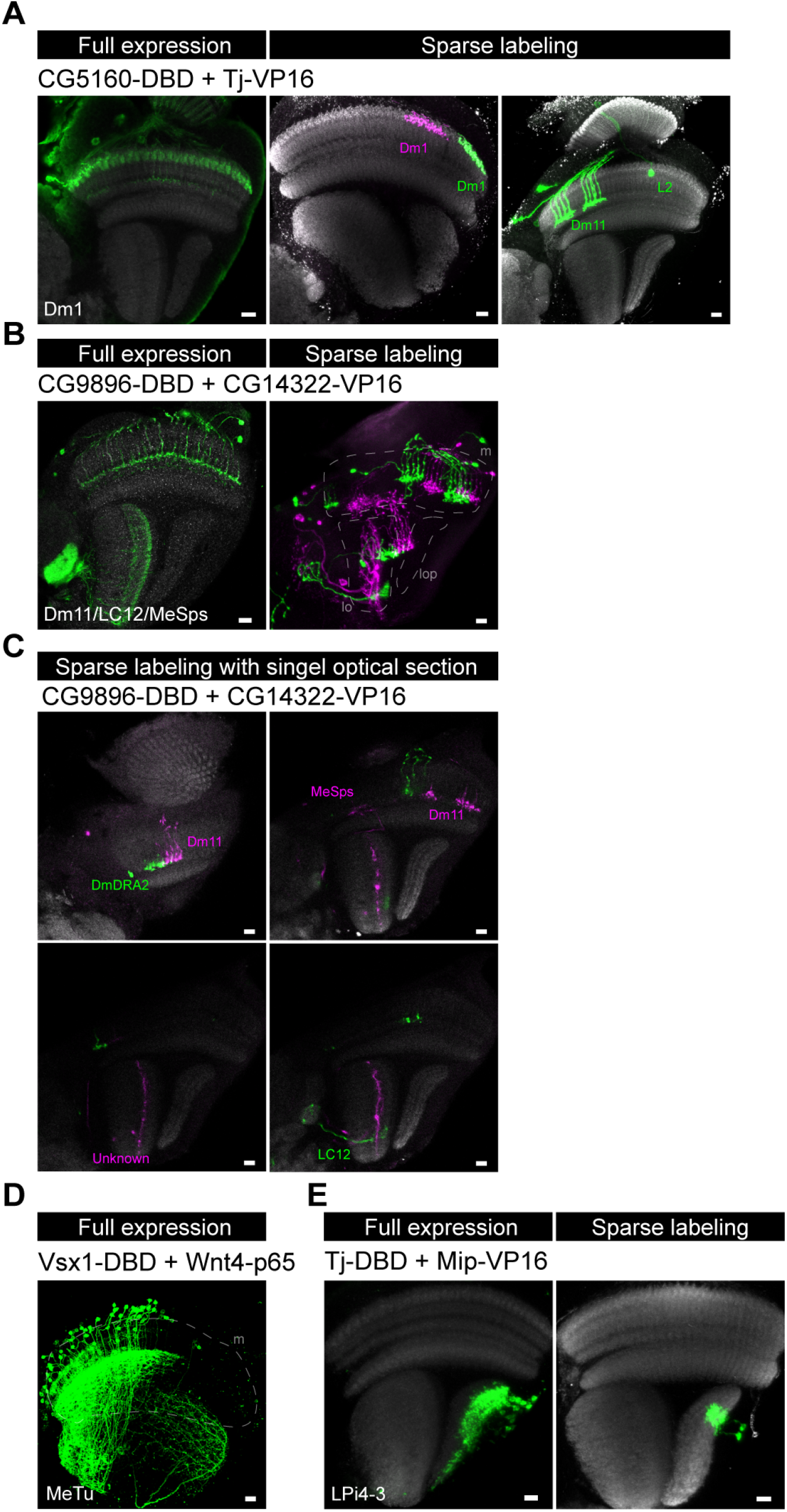
Characterization of selected gene-specific split-GAL4 lines targeting different cell types/clusters. Targeted cell types predicted by scRNAseq expression are shown in the lower left corner for each split-GAL4 line. The expression pattern of each split-GAL4 line is shown either with UAS-myr-GFP reporter for full expression (right) or with UAS-MCFO lines for sparse labeling (right). (A) A different split-GAL4 combination targeting Dm1. We also observed Dm11 and L2 neurons in this line. (B-C) A different split-GAL4 combination targeting Dm11, LC12, and MeSps neurons. Sparse labeling confirmed the presence of all three cell types. In addition, there is one additional unknown cell type labeled in the lobula. (D) The cell bodies of MeTu neurons are located in the dorsal half of medulla cortex in adult. (E) A different split-GAL4 combination targeting LPi4-3. Anti-NCad staining (gray) is used for visualizing neuropils. Images are substack projections from full expression labeling or segmented single cells from sparse labeling to show distinct morphological features of distinct cell types. Scale bar: 10 *μ*m.

**Fig. S2.**
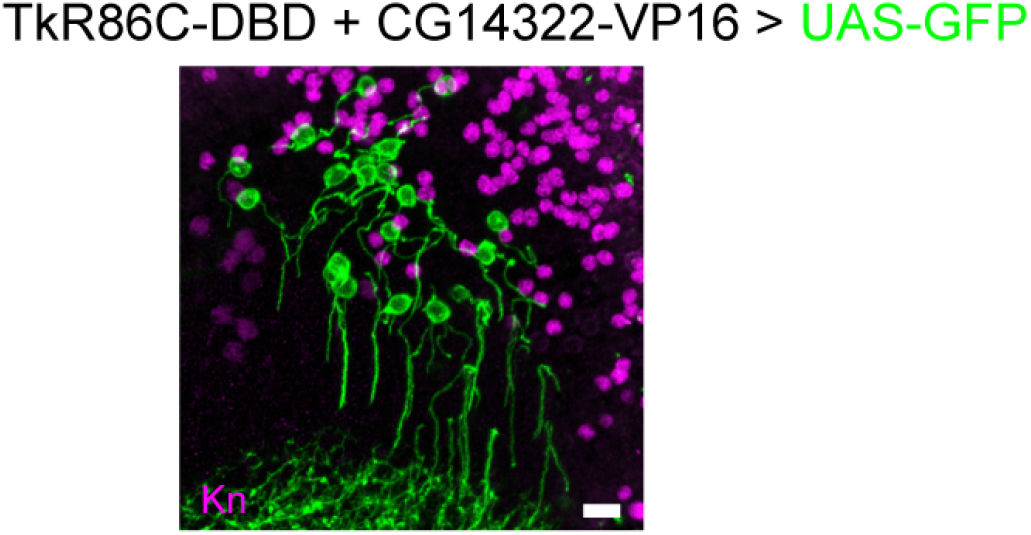
Co-staining of MeSps neurons labeled by TkR86C-DBD and CG14322-VP16 with anti-Kn (Magenta). MeSps neurons expressing GFP reporter are Kn negative. Scale bar: 10 *μ*m

**Fig. S3.**
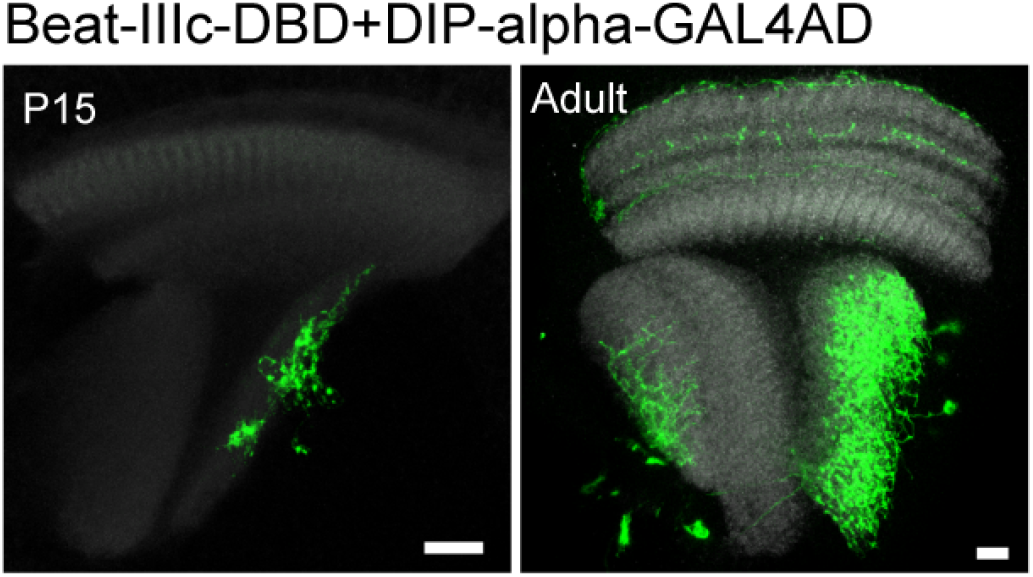
The full expression pattern of Beat-IIIc-GAL4DBD + DIP-GAL4AD line is shown with UAS-myr-GFP reporter at P15 (left) and adult (right) stages. Note that the targeted cell type (LPi4-3) is always observed at multiple developmental stages, although other cell types might be observed in addition to it. Anti-NCad staining (gray) is used for visualizing neuropils. Images are substack projections from full expression labeling to show distinct morphological features of distinct cell types. Scale bar: 10 *μ*m.

**Fig. S4.**
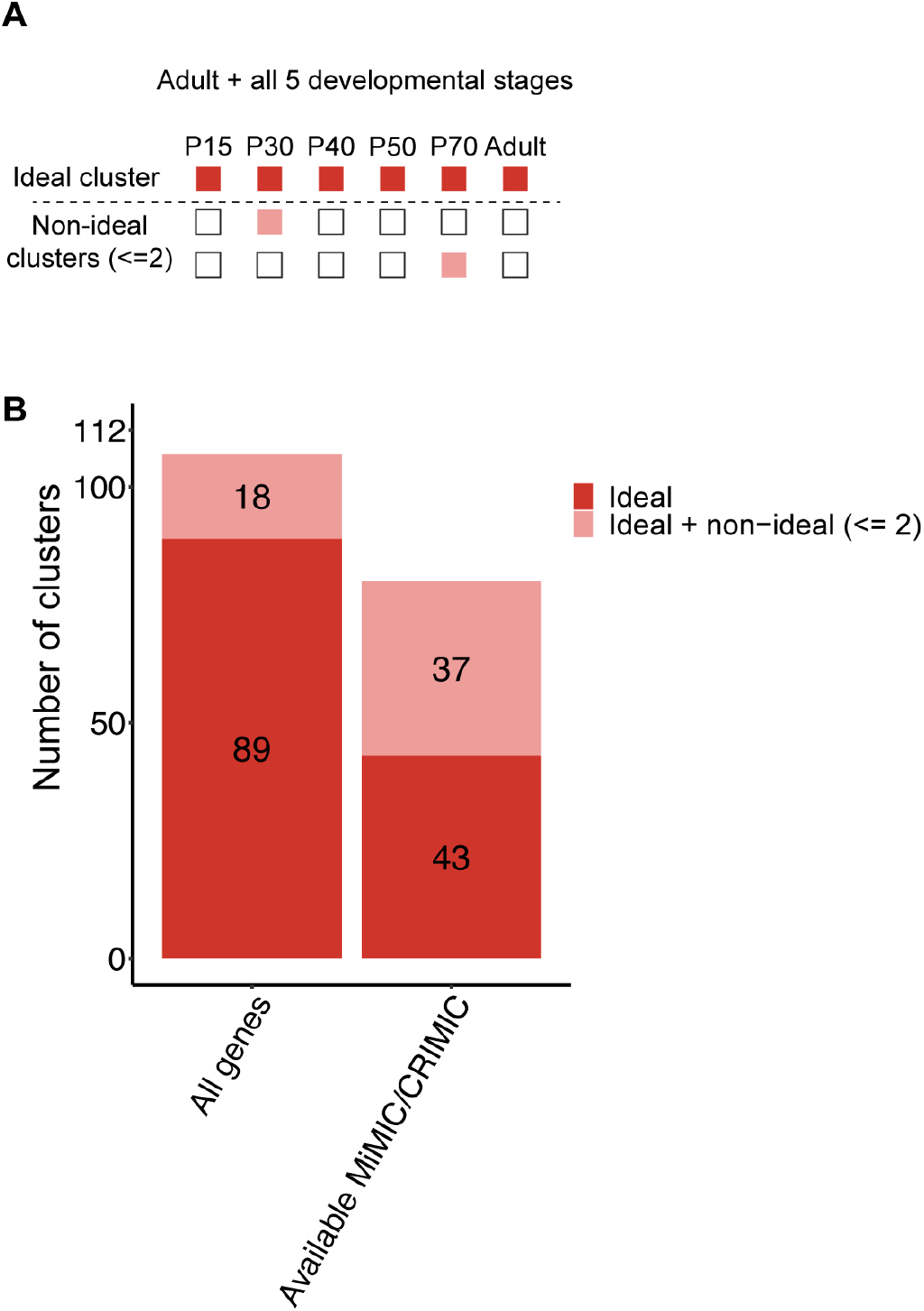
(A) Illustration of the strategy used to identify gene pairs that mark a previously unannotated cluster of interest. Red boxes indicate the stages when a gene pair is predicted to be on in the cluster of interest while pink boxes indicate when the gene pair is predicted to be active in other clusters. (B) Number of previously unannotated clusters that are predicted to be identified with gene pairs when all genes detected in the atlas are considered (left) or when only genes with coding intronic MiMIC or CRIMIC lines available are considered (right).

**Fig. S5.**
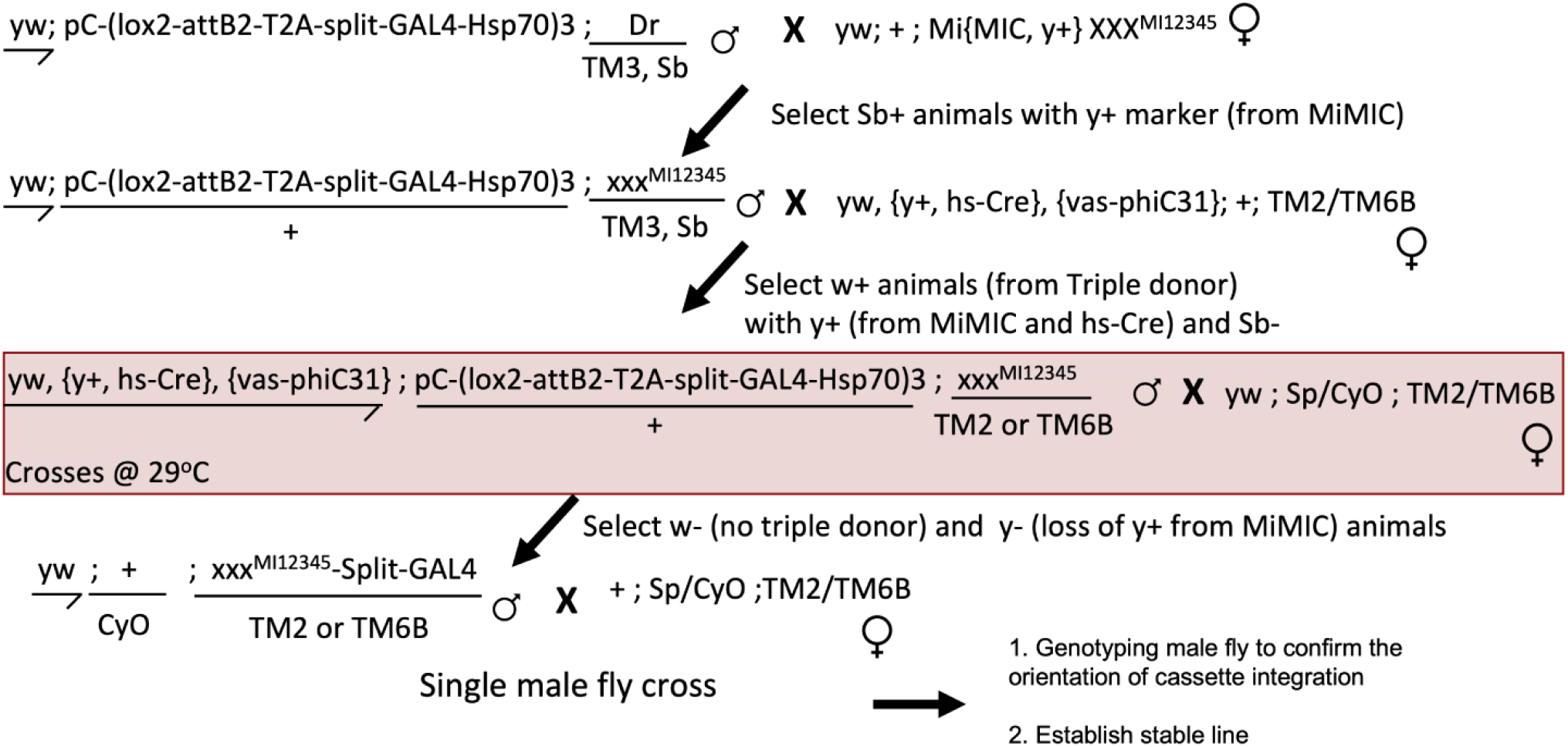
*in vivo* genetic crossing scheme for swapping T2A-split GAL4 in coding intronic MiMIC lines. Flies with hs-Cre, vas-C31, split-GAL4 donor and targeting MiMIC cross should be incubated at 29°C to induce the expression of Cre recombination.

## Notes

### Competing Interest Statement

The authors have declared no competing interest.

## References

1. AH Brand, N Perrimon, Targeted gene expression as a means of altering cell fates and generating dominant phenotypes. Dev. (Cambridge, England) 118, 401–415 (1993).

2. SL Lai, T Lee, Genetic mosaic with dual binary transcriptional systems in drosophila. Nat. Neurosci. 9, 703–709 (2006).

3. CJ Potter, B Tasic, EV Russler, L Liang, L Luo, The q system: a repressible binary system for transgene expression, lineage tracing, and mosaic analysis. Cell 141, 536–548 (2010).

4. A Jenett, et al., A GAL4-driver line resource for drosophila neurobiology. Cell Reports 2, 991–1001 (2012).

5. EZ Kvon, et al., Genome-scale functional characterization of drosophila developmental enhancers in vivo. Nature 512, 91–95 (2014).

6. H Luan, F Diao, RL Scott, BH White, The drosophila split gal4 system for neural circuit mapping. Front. Neural Circuits 14, 603397 (2020).

7. H Luan, NC Peabody, CR Vinson, BH White, Refined spatial manipulation of neuronal function by combinatorial restriction of transgene expression. Neuron 52, 425–436 (2006).

8. BD Pfeiffer, et al., Refinement of tools for targeted gene expression in drosophila. Genetics 186, 735–755 (2010).

9. CY Ting, et al., Focusing transgene expression in drosophila by coupling gal4 with a novel split-LexA expression system. Genetics 188, 229–233 (2011).

10. H Dionne, KL Hibbard, A Cavallaro, JC Kao, GM Rubin, Genetic reagents for making split-GAL4 lines in drosophila. Genetics 209, 31–35 (2018) Publisher: Oxford University Press.

11. L Tirian, BJ Dickson, The VT GAL4, LexA, and split-GAL4 driver line collections for targeted expression in the drosophila nervous system. BioRxiv p. 198648 (2017) Publisher: Cold Spring Harbor Laboratory.

12. GW Meissner, et al., A searchable image resource of <em>drosophila</em> GAL4-driver expression patterns with single neuron resolution. bioRxiv p. 2020.05.29.080473 (2022).

13. H Otsuna, M Ito, T Kawase, Color depth MIP mask search: a new tool to expedite split-GAL4 creation. bioRxiv p. 318006 (2018).

14. YZ Kurmangaliyev, J Yoo, J Valdes-Aleman, P Sanfilippo, SL Zipursky, Transcriptional programs of circuit assembly in the drosophila visual system. Neuron 108, 1045–1057.e6 (2020-12-23).

15. MN Özel, et al., Neuronal diversity and convergence in a visual system developmental atlas. Nature 589, 88–95 (2021).

16. JR Bateman, AM Lee, Ct Wu, Site-specific transformation of drosophila via phiC31 integrase-mediated cassette exchange. Genetics 173, 769–777 (2006).

17. F Diao, et al., Plug-and-play genetic access to drosophila cell types using exchangeable exon cassettes. Cell Reports 10, 1410–1421 (2015).

18. PT Lee, et al., A gene-specific t2a-GAL4 library for drosophila. eLife 7, e35574 (2018).

19. KJT Venken, et al., MiMIC: a highly versatile transposon insertion resource for engineering drosophila melanogaster genes. Nat. Methods 8, 737–743 (2011).

20. FP Davis, et al., A genetic, genomic, and computational resource for exploring neural circuit function. eLife 9, e50901 (2020) Publisher: eLife Sciences Publications, Ltd.

21. A Nern, BD Pfeiffer, GM Rubin, Optimized tools for multicolor stochastic labeling reveal diverse stereotyped cell arrangements in the fly visual system. Proc. Natl. Acad. Sci. 112, E2967–E2976 (2015) Publisher: Proceedings of the National Academy of Sciences.

22. JC Tuthill, A Nern, SL Holtz, GM Rubin, MB Reiser, Contributions of the 12 neuron classes in the fly lamina to motion vision. Neuron 79, 128–140 (2013).

23. M Wu, et al., Visual projection neurons in the drosophila lobula link feature detection to distinct behavioral programs. eLife 5, e21022 (2016).

24. LS Gramates, et al., Fly base: a guided tour of highlighted features. Genetics 220, iyac035 (2022).

25. Q Duan, R Estrella, A Carson, Y Chen, PC Volkan, <em>drosophila attP40</em> back-ground alters glomerular organization of the olfactory receptor neuron terminals. bioRxiv p. 2022.06.16.496338 (2022).

26. CM Groen, JL Podratz, J Pathoulas, N Staff, AJ Windebank, Genetic reduction of mitochondria complex i subunits is protective against cisplatin-induced neurotoxicity in drosophila. The J. Neurosci. The Off. J. Soc. for Neurosci. 42, 922–937 (2022).

27. K van der Graaf, S Srivastav, P Singh, JA McNew, M Stern, The drosophila <em>attP40</em> docking site and derivatives are insertion mutations of <em>MSP300</em>. bioRxiv p. 2022.05.14.491875 (2022).

28. Stan Development Team, The Stan Core Library (2022).

29. R Core Team, R: A language and environment for statistical computing (2022).

30. L Scrucca, M Fop, TB Murphy, AE Raftery, mclust 5: clustering, classification and density estimation using Gaussian finite mixture models. The R J. 8, 289–317 (2016).

