## Supplementary material for "Using single-cell RNA sequencing to generate cell-type-specific split-GAL4 reagents throughout development": Link to the Materials and Methods

**Supplementary Table 1: Fly strains and antibodies**

| REAGENT or RESOURCE | SOURCE | IDENTIFIER |
|---|---|---|
| Fly strains | | |
| Drosophila, yw;; | Desplan lab | N/A |
| Drosophila, 10XUAS-myr-GFP | BDSC | BDSC #32197, #32198 |
| Drosophila, R57C10-Flp2::PEST; 10xUAS(FRT.stop)myr::smGdP-V5-THS-10xUAS(FRT.stop)myr::smGdP-FLAG} ; GMR-myr::RFP, Rh4-lacZ | Desplan lab and BDSC | BDSC #62124 |
| Drosophila, 5-HT1B[MI05213]-T2A-GAL4DBD | This study | N/A |
| Drosophila, 5-HT2B[MI06500]-T2A-GAL4DBD | This study | N/A |
| Drosophila, ab-T2A-GAL4DBD | This study | N/A |
| Drosophila, AdoR[MI01202]-T2A-GAL4DBD | This study | N/A |
| Drosophila, ara-T2A-GAL4DBD | This study | N/A |
| Drosophila, Acj6[MI07818]-T2A-GAL4DBD | This study | N/A |
| Drosophila, BBS4[MI15481]-T2A-GAL4DBD | This study | N/A |
| Drosophila, Beat-IIIc[MI03726]-T2A-GAL4DBD | This study | N/A |
| Drosophila, bi-T2A-GAL4DBD | This study | N/A |
| Drosophila, bt[MI06578]-T2A-GAL4DBD | This study | N/A |

| | | |
|---|---|---|
| Drosophila, CG11317[MI04510]-T2A-GAL4DBD | This study | N/A |
| Drosophila, CG11537[MI04840]-T2A-GAL4DBD | This study | N/A |
| Drosophila, CG13375[MI15214]-T2A-GAL4DBD | This study | N/A |
| Drosophila, CG13408[CR02202]-T2A-GAL4DBD | This study | N/A |
| Drosophila, CG14322[MI11688]-T2A-GAL4DBD | This study | N/A |
| Drosophila, CG14521(DIPγ)[MI03222]-T2A-GAL4DBD | (1) | N/A |
| Drosophila, CG15353-T2A-GAL4DBD | This study | N/A |
| Drosophila, CG1688[CR01570]-T2A-GAL4DBD | This study | N/A |
| Drosophila, CG2772[CR02306]-T2A-GAL4DBD | This study | N/A |
| Drosophila, CG31183[MI02001]-T2A-GAL4DBD | This study | N/A |
| Drosophila, CG31689[MI03245]-T2A-GAL4DBD | This study | N/A |
| Drosophila, CG32432[MI03670]-T2A-GAL4DBD | This study | N/A |
| Drosophila, CG34411[MI06794]-T2A-GAL4DBD | This study | N/A |
| Drosophila, CG42368[MI09901]-T2A-GAL4DBD | This study | N/A |

| | | |
|---|---|---|
| Drosophila, CG43737[MI10158]-T2A-GAL4DBD | This study | N/A |
| Drosophila, CG44325[MI09901]-T2A-GAL4DBD | This study | N/A |
| Drosophila, CG5160-T2A-GAL4DBD | This study | N/A |
| Drosophila, CG5910[MI09800]-T2A-GAL4DBD | This study | N/A |
| Drosophila, CG7191[MI03423]-T2A-GAL4DBD | This study | N/A |
| Drosophila, CG9896-T2A-GAL4DBD | This study | N/A |
| Drosophila, CngA[MI05524]-T2A-GAL4DBD | This study | N/A |
| Drosophila, CNMaR[MI01128]-T2A-GAL4DB | This study | N/A |
| Drosophila, Fer2[MI09483]-T2A-GAL4DBD | This study | N/A |
| Drosophila, foxo[MI00493]-T2A-GAL4DBD | This study | N/A |
| Drosophila, glob1[MI14203]-T2A-GAL4DBD | This study | N/A |
| Drosophila, heph[MI04030]-T2A-GAL4DBD | This study | N/A |
| Drosophila, hig[MI05774]-T2A-GAL4DBD | This study | N/A |
| Drosophila, Kek3[CR00930]-T2A-GAL4DBD | This study | N/A |
| Drosophila, Lkr[MI08640]-T2A-GAL4DBD | This study | N/A |

| | | |
|---|---|---|
| Drosophila, Meltrin[MI09878]-T2A-GAL4DBD | This study | N/A |
| Drosophila, mld[MI07175]-T2A-GAL4DBD | This study | N/A |
| Drosophila, ort-T2A-GAL4DBD | This study | N/A |
| Drosophila, PHDP[CR01516]-T2A-GAL4DBD | This study | N/A |
| Drosophila, Pka-R1[MI06935]-T2A-GAL4DBD | This study | N/A |
| Drosophila, rdo[MI14128]-T2A-GAL4DBD | This study | N/A |
| Drosophila, Reck-T2A-GAL4DBD | This study | N/A |
| Drosophila, rst[MI04842]-T2A-GAL4DBD | This study | N/A |
| Drosophila, RunxB[MI10372]-T2A-GAL4DBD | This study | N/A |
| Drosophila, Rya-R[MI01843]-T2A-GAL4DBD | This study | N/A |
| Drosophila, salm[MI13018]-T2A-GAL4DBD | This study | N/A |
| Drosophila, Tj-T2A-GAL4DBD | This study | N/A |
| Drosophila, TkR86C[MI05788]-T2A-GAL4DBD | This study | N/A |
| Drosophila, TrissinR[MI05488]-T2A-GAL4DBD | This study | N/A |
| Drosophila, tsh-T2A-GAL4DBD | This study | N/A |
| Drosophila, Tusp[MI04698] -T2A-GAL4DBD | This study | N/A |

| | | |
|---|---|---|
| Drosophila, Vsx1-T2A-GAL4DBD | This study | N/A |
| Drosophila, Wnt10-T2A-GAL4DBD | (2) | N/A |
| Drosophila, Beat-IIIc[MI03726]-T2A-VP16 | This study | N/A |
| Drosophila, bru1[MI00135]-T2A-VP16 | This study | N/A |
| Drosophila, CCAP-R[MI05804]-T2A-VP16 | This study | N/A |
| Drosophila, CG14010 (DIPη)[MI07948]-T2A-VP16 | This study | N/A |
| Drosophila, CG14322[MI11688]-T2A-VP16 | This study | N/A |
| Drosophila, CG14431[MI10741]-T2A-VP16 | This study | N/A |
| Drosophila, CG14521(DIPγ)[MI03222]-T2A-VP16 | (1) | N/A |
| Drosophila, CG14947-T2A-VP16 | This study | N/A |
| Drosophila, CG2016[MI04995]-T2A-p65 | This study | N/A |
| Drosophila, CG31191[MI07868]-T2A-VP16 | This study | N/A |
| Drosophila, CG31689[MI03245]-T2A-VP16 | This study | N/A |
| Drosophila, CG32105-T2A-VP16 | This study | N/A |
| Drosophila, CG4238[MI13092]-T2A-VP16 | This study | N/A |

| | | |
|---|---|---|
| Drosophila, CG43737[MI10158]-T2A-VP16 | This study | N/A |
| Drosophila, CG5397[MI15629]-T2A-VP16 | This study | N/A |
| Drosophila, CG9109[MI01660]-T2A-VP16 | This study | N/A |
| Drosophila, CG9743[MI09362]-T2A-VP16 | This study | N/A |
| Drosophila, dac-T2A-GAL4AD | This study | N/A |
| Drosophila, Dop1R2[CR02529]-T2A-VP16 | This study | N/A |
| Drosophila, erm-T2A-VP16 | This study | N/A |
| Drosophila, eya-T2A-VP16 | This study | N/A |
| Drosophila, Fife[MI08414]-T2A-VP16 | This study | N/A |
| Drosophila, fs[MI04308]-T2A-VP16 | This study | N/A |
| Drosophila, Gad1[MI09277]-T2A-VP16 | (3) | BDSC #60322 |
| Drosophila, Kn[MI15480]-T2A-p65 | (4) | N/A |
| Drosophila, Lim1-T2A-VP16 | This study | N/A |
| Drosophila, mbl[MI00139]-T2A-VP16 | This study | N/A |
| Drosophila, Mip-T2A-VP16 | This study | N/A |
| Drosophila, mirr-T2A-VP16 | This study | N/A |
| Drosophila, MsR2[CR01840]-T2A-p65 | This study | N/A |
| Drosophila, NetA[MI04563]-T2A-VP16 | This study | N/A |

| | | |
|---|---|---|
| Drosophila, orb[MI04761]-T2A-VP16 | This study | N/A |
| Drosophila, ort-T2A-VP16 | This study | N/A |
| Drosophila, Ple-T2A-p65 | (5) | BDSC #84712 |
| Drosophila, rn[MI09946]-T2A-VP16 | This study | N/A |
| Drosophila, run-T2A-VP16 | This study | N/A |
| Drosophila, Rx[CR00377]-T2A-p65 | This study | N/A |
| Drosophila, sfl[MI02467]-T2A-p65 | This study | N/A |
| Drosophila, slou-T2A-VP16 | This study | N/A |
| Drosophila, SoxN-T2A-VP16 | This study | N/A |
| Drosophila, svp[MI01102]-T2A-VP16 | This study | N/A |
| Drosophila, tey-T2A-VP16 | This study | N/A |
| Drosophila, tj-T2A-VP16 | This study | N/A |
| Drosophila, tsh-T2A-VP16 | This study | N/A |
| Drosophila, VGlut[MI04979]-T2A-p65 | (6) | BDSC #82986 |
| Drosophila, Wnt10-T2A-p65 | (2) | N/A |
| Drosophila, P{lox(Trojan-GAL4DBD)x3} | This study | N/A |
| Drosophila, P{lox(Trojan-VP16)x3} | This study | N/A |
| Drosophila, P{loxP(Trojan-GAL4DBD.0)} | This study | N/A |
| Drosophila, P{loxP(Trojan-GAL4DBD.1)} | This study | N/A |

| | | |
|---|---|---|
| Drosophila, P{loxP(Trojan-GAL4DBD.2)} | This study | N/A |
| Drosophila, P{loxP(Trojan-VP16.0)} | This study | N/A |
| Drosophila, P{loxP(Trojan-VP16.1)} | This study | N/A |
| Drosophila, P{loxP(Trojan-VP16.2)} | This study | N/A |
| Antibodies | | |
| Sheep anti-GFP (1:200) | BioRad | 4745-1051 (RRID:AB_619712) |
| Chicken anti-V5 (1:5000) | Abcam | ab9113 (RRID:AB_307022) |
| Rat anti-FLAG (1:200) | Novus Biologicals | NBP1-06712 (RRID:AB_1625981) |
| Rat anti-NCad (1:20) | DSHB | DN-ex#8 (RRID:AB_528121) |
| Mouse anti-Chaoptin (1:20) | DSHB | 24B10 (RRID:AB_528161) |
| Mouse anti-Brp (1:20) | DSHB | nc82 (RRID:AB_2314866) |
| Mouse anti-pros (1:33) | DSHB | MR1A (RRID:AB_528440) |
| Guinea pig anti-Tj (1:250) | (7), Gift from Dorothea Godt | N/A |
| Rabbit anti-Toy (1:500) | (8), Desplan lab | N/A |
| Guinea pig anti-Fd59A (1:400) | (9), Gift from James Skeath | N/A |
| Guinea pig anti-Kn (1:200) | (10), Desplan lab | N/A |
| Donkey anti-sheep Alexa Fluor 488 (1:500) | Jackson ImmunoResearch | 713-545-147 (RRID:AB_2340745) |

| | | |
|---|---|---|
| Donkey anti-chicken Alexa Fluor 488 (1:500) | Jackson ImmunoResearch | 703-545-155 (RRID:AB_2340375) |
| Donkey anti-rat Cy3 (1:500) | Jackson ImmunoResearch | 712-165-153 (RRID:AB_2340667) |
| Donkey anti-mouse Alexa Fluor 555 (1:500) | Invitrogen | A31570 (RRID:AB_2536180) |
| Donkey anti-rabbit Alexa Fluor 555 (1:500) | Invitrogen | A31572 (RRID:AB_162543) |
| Donkey anti-rat Alexa Fluor 647 (1:200) | Jackson ImmunoResearch | 712-605-153 (RRID:AB_2340694) |
| Donkey anti-guinea pig Alexa Fluor 647 (1:200) | Jackson ImmunoResearch | 706-605-148 (RRID:AB_2340476) |

**Referneces**


1. M. Courgeon, C. Desplan, Coordination between stochastic and deterministic specification in the Drosophila visual system. *Science* **366**, eaay6727 (2019).

2. B. Ewen-Campen, T. Comyn, E. Vogt, N. Perrimon, No Evidence that Wnt Ligands Are Required for Planar Cell Polarity in Drosophila. *Cell Rep* **32**, 108121 (2020).

3. F. Diao, *et al*., Plug-and-play genetic access to drosophila cell types using exchangeable exon cassettes. *Cell Rep* **10**, 1410–1421 (2015).

4. Q. Xie, *et al*., Temporal evolution of single-cell transcriptomes of Drosophila olfactory projection neurons. *Elife* **10**, e63450 (2021).

5. B. Deng, *et al*., Chemoconnectomics: Mapping Chemical Transmission in Drosophila. *Neuron* **101**, 876-893.e4 (2019).

6. H. Lacin, *et al*., Neurotransmitter identity is acquired in a lineage-restricted manner in the Drosophila CNS. *Elife* **8**, e43701 (2019).

7. F. Gunawan, M. Arandjelovic, D. Godt, The Maf factor Traffic jam both enables and inhibits collective cell migration in Drosophila oogenesis. *Development* **140**, 2808–2817 (2013).

8. M. N. Özel, *et al*., Neuronal diversity and convergence in a visual system developmental atlas. *Nature* **589**, 88–95 (2021).

9. H. Lacin, *et al.*, Genome-wide identification of Drosophila Hb9 targets reveals a pivotal role in directing the transcriptome within eight neuronal lineages, including activation of nitric oxide synthase and Fd59a/Fox-D. *Dev Biol* **388**, 117–133 (2014).

10. N. Konstantinides, *et al.*, A complete temporal transcription factor series in the fly visual system. *Nature* **604**, 316–322 (2022).
